# Simultaneous enhancement of multiple stability properties using two-parameter control methods in *Drosophila melanogaster*

**DOI:** 10.1101/027169

**Authors:** Sudipta Tung, Abhishek Mishra, Sutirth Dey

**Author notes:** Correspondence to Name and address of the corresponding author: Sutirth Dey Biology Division Indian Institute of Science Education and Research Pune Dr. Homi Bhabha Road,Pune, Maharashtra, India 411 008.

## Abstract

Although a large number of methods have been proposed to control the non-linear dynamics of unstable popuations, very few have been actually adopted for application. One reason for this gap is the fact that few control methods have been empirically verified using biological populations. To address this issue, we investigated the effects of two well-studied control methods (Both Limiter Control and Target-Oriented Control) on the dynamics of unstable populations of *Drosophila melanogaster*. We show that both methods can significantly reduce population fluctuations, decrease extinction probability and increase effective population size simultaneously. This is in contrast with single parameter control methods that are not able to achieve multiple aspects of stability at the same time. We use the distribution of population sizes to derive biologically intuitive explanations for the mechanisms of how these two control methods attain stability. Finally, we show that non-Drosophila specific biologically realistic simulations are able to capture almost all the trends of our data. This shows that our results are likely generalizable over a wide range of taxa. The primary insight of our study is that control methods that incorporate both culling and restocking have better all-round performance in terms of stabilizing populations.

## 1. INTRODUCTION

Although several methods have been proposed for stabilizing biological populations over the last two decades (e.g. Dattani et al. 2011; Güémez and Matías 1993; Hilker and Westerhoff 2005; McCallum 1992; Sah et al. 2013), few (if any) have been actually applied for conserving threatened populations. This gap between theory and application has several putative reasons. Firstly, control methods are often investigated in the context of simple 1-D models (McCallum 1992) which are typically not good descriptors of the dynamics of “charismatic species” like mammals or birds. Unfortunately, it is the charismatic species that are generally the focus of most conservation efforts (Kontoleon and Swanson 2003) and therefore most practising conservation biologists find little use for much of the proposed control methods. However, it should be noted here that simple 1-D models are often found to be good descriptors of the dynamics of a large number of non-charismatic species. This is not surprising since the derivations of many of these models are based on assumptions about the kind of competition and spatial distribution of organisms that are valid over a wide range of taxa (Brännström and Sumpter 2005). For example, the Ricker model has been shown to be a good descriptor of the dynamics of, *inter alia*, bacteria (Ponciano et al. 2005), fungi (Ives et al. 2004), ciliates (Fryxell et al. 2005), insects (Sheeba and Joshi 1998) and fishes (Denney et al. 2002). Together, these organisms represent an overwhelming majority of the biodiversity found on this planet, at least a fraction of which has already been recorded to be extinct (Baillie and Butcher 2012). Thus, the conservation of such non-charismatic species is likely to become important sooner than later and it is essential to investigate the methods to conserve them.

Another major issue with several existing theoretical studies lies in the plurality of the concept of stability. In ecology, the term stability can refer to many concepts (Grimm and Wissel 1997) and most theoretical studies typically restrict to any one of them. However, in any real world usage, the adopted control method must be able to simultaneously stabilize multiple aspects of the dynamics. Thus, for example, a method that reduces fluctuations in population sizes, but has relatively less impact on extinction probability, is of limited utility. Since different aspects of often do not correlate with each other (Dey and Joshi 2013; Dey et al. 2008), choosing a method often becomes problematic.

To complicate matters further, most control methods proposed in the theoretical literature lack adequate empirical (i.e. ones that use real biological populations as opposed to computer simulations) verification. Some of the most well-known empirical studies on population control deal with methods that either require high levels of mathematical sophistication (e.g. Desharnais et al. 2001) or very detailed models of the system (Becks et al. 2005). The high degree of specificity of these studies can sometimes make it difficult to extend their insights into controlling other systems. Moreover, such studies (Becks et al. 2005; Desharnais et al. 2001) often deal with amelioration of chaos or characterization of the attractor, whereas the primary concern of most conservation biologists would be to prevent inbreeding or reduce extinction probability. Consequently, such empirically well-characterized control methods turn out to be of limited relevance for most real-world applications. As stated already, much of the proposed control methods have never been investigated using biological populations. Given that the survivals of species are at stake, the reluctance of the practitioners in adopting untested methods for controlling natural populations is well justified. The only way to bridge this gap between theory and practise is to empirically verify the control methods proposed in the literature under conditions that are as close to their conditions of intended use as possible. Clearly, methods that require relatively less system-specific information and are easier to implement (e.g. Gusset et al. 2009; McCallum 1992), are likely to be more useful in this context. One such set of control methods is the so-called limiter class of methods.

Broadly speaking, the limiter methods do not allow the populations to go above or below (depending on the method) a pre-determined threshold through culling or restocking respectively. Recent empirical studies have shown that such methods can typically reduce either fluctuations in population sizes or extinction probability, but not both (Sah et al. 2013; Tung et al. 2015). This observation led to the conjecture that methods which involve both restocking and culling might prove to be more successful in simultaneous control of multiple aspects of stability. A well-studied method of this type is the Target-Oriented Control (TOC) (Braverman and Chan 2014; Braverman and Franco 2015; Dattani et al. 2011; Franco and Liz 2013) which is a modification of the traditional proportional feedback method (Güémez and Matías 1993; Liz 2010). In TOC, the current population size is perturbed towards a predetermined ‘target’ by culling or restocking (Dattani et al. 2011; Franco and Liz 2013). The magnitude of the perturbation is determined based on the difference between the pre-perturbation population size and the target value. Theoretical studies have shown that TOC globally stabilizes higher order difference equations (Braverman and Franco 2015) and is particularly useful in those cases where the population size needs to be manipulated towards a pre-determined value (Dattani et al. 2011).

Another method that involves both culling and restocking is the recently proposed Both Limiter Control or BLC, which involves setting an upper and a lower threshold *a priori* (Tung et al.2014). Each time the population size is outside the range set by these thresholds, appropriate culling or restocking is implemented to bring the size back to the upper or the lower threshold respectively. It has been shown numerically that BLC can protect populations from overcrowding and extinction risk due to demographic stochasticity (Tung et al. 2014). However, till date, there has been no empirical investigation of how these two control methods affect the dynamics of real biological populations.

In this study, we investigate the effects of BLC and TOC in stabilizing the dynamics of spatially-unstructured laboratory populations of *Drosophila melanogaster*. Both these methods were found to be capable of inducing significant reduction in fluctuations in population sizes and extinction probability. Moreover, both methods also significantly increased the effective population sizes. However, the superior performance of BLC and TOC came at the cost of a significantly large effort magnitude, which is likely to translate into relatively high economic expenditure. We also derive biologically intuitive understandings of how the control methods work by comparing the distribution of population sizes with and without control. Finally, we show that simulations using a biologically realistic, non-*Drosophila*-specific model, can capture most of the trends of our experimental results. This suggests that our observations are likely to be generalizable over a wide range of taxonomic groups.

## 2. METHODS

### 2.1 Experimental populations and their maintenance

In this study, we used 20 single vial populations of *Drosophila melanogaster* derived from a large outbred population, called DB_4_. The ancestry and maintenance regime of the outbred population is mentioned elsewhere (Sah et al. 2013). Each single vial culture was initiated by collecting 10 eggs on 1.1 ml of standard banana-jaggery medium and reared in an incubator at 25? C temperature under constant light. Once the adults started eclosing in a vial, they were transferred daily into corresponding adult-holding vials containing ~6ml of medium. The correspondence between an egg-collection vial and its adult-holding vial was strictly maintained. On the 18th day from the day of egg collection, the flies were supplied with live yeast paste to boost fecundity. Three days later, the adults were counted under mild CO_2_ anaesthesia and culling or restocking were implemented wherever necessary as per the protocol of the corresponding control method (see section 2.2). After this, the adults were put into fresh vials containing 1.1 ml food for oviposition. After 24 hours, the adults were discarded and the eggs formed the next generation. The experiment was 14 generation long and all indices (except extinction probability) were computed on the time series of the breeding population (i.e. after the implementation of restocking/culling).

### 2.2 Control methods

*Both Limiter Control (BLC):* In BLC, the population size is not allowed to go beyond predetermined upper and lower threshold values (Sah et al. 2013). Mathematically, this can be represented as *N*_*t*_′ *= max(min(N_t_, U), L)*, where *N*_*t*_ and *N*_*t*_*′* are population sizes before and after the application of the control method, *U* and *L* are the pre-determined values of the upper and lower thresholds, and *max* and *min* denote the maximum and minimum operators. Here, we arbitrarily chose the lower and upper thresholds as 4 and 10 respectively. Since the dynamics of sexually reproducing species are primarily driven by the number of females in the population, we restricted the implementation of the control to the females. In other words, when the number of females in a given generation was less than 4 or more than 10, BLC was applied by restocking to 4 females or culling to 10 females respectively.

*Target Oriented Control (TOC)*: In TOC, the population size is nudged towards an *a priori* fixed target value (Dattani et al. 2011). It is a two-parameter control method which can be mathematically represented as *N*_*t*_′ = *N*_*t*_ + *c*_*d*_ × (*Ʈ* -*N*_*t*_), where *N*_*t*_ and *N*_*t*_*′* are population sizes before and after the application of TOC and *Ʈ* denotes the target population size. The parameter *c*_*d*_ (arbitrarily set to 0.7 here) represents the fraction by which the difference between the target and current population size is restocked or culled. Thus, in our experiment, when the population size exceeds the target (*Ʈ*), 70% of the excess individuals are culled and when population size is below the target, 70%of the difference in number of individuals is added to the population. Simulations suggest that when the target is kept close to the carrying capacity of the unperturbed population, TOC requires very little external perturbation in the long term (Dattani et al. 2011). To obtain an estimate of the carrying capacity (*K*), we fitted the Ricker model (Ricker 1954) to time series data of unperturbed *Drosophila* populations, under similar maintenance regime, from one of our earlier studies (Sah et al. 2013). This led to a mean value of 25 ± 8.4 (standard deviation). Since the Ricker model shows stable-point equilibrium at *K*, we arbitrarily set the value of the target at 30. This slight departure from the estimated value was implemented because precise estimates of *K* are unlikely to be obtained for most real scenarios.

Since earlier theoretical studies (Dattani et al. 2011; Tung et al. 2014) had suggested TOC to be a very effective control method, we decided to test it under somewhat more stringent conditions than BLC. For this, we incorporated some degree of imprecision in the implementation of the control method. Modifying the protocol of an earlier study (Dey and Joshi 2006), we estimated the number of females to be added or removed as *floor* [0.5 × *c*_*d*_ × (*Ʈ*-*N*_*t*_)] where *floor [x]* denotes the function leading to the largest integer not greater than x. This way of calculating the magnitude of the control assumes an equal sex ratio which will not be the case in every generation, thus introducing some noise in the implementation.

### 2.3 Unperturbed populations and experimental replicates

Apart from the BLC and TOC lines, we also had two sets of populations that were kept unperturbed, and maintained as mentioned in section 2.1. These two sets were called C1 and C2, and they served as unperturbed reference populations for BLC and TOC respectively. This pairing was done at the time of the experimental set up. The dynamics of the C1 and C2 populations have already been reported previously (Tung et al. 2015). When the C1 or C2 populations went extinct, they were reset by adding 4 males and 4 females from outside, which contributed to the effort magnitude (see below) of these populations. There were no needs for separate resets in the BLC and TOC lines, since the control methods automatically ensured that the extinct populations were rescued. All flies that were used for restocking were maintained as mentioned in section 2.1 except for the fact that they were provided with ~5 ml of larval food.Overall, there were four treatments (BLC, TOC, C1 and C2) in this study and each consisted of 5 replicate populations.

### 2.4 Stability measures

We used three measures of stability, namely constancy, persistence and effective population size. Constancy stability (Grimm and Wissel 1997) of populations refers to the magnitude of temporal fluctuations in population sizes: population that have larger fluctuations have lesser constancy stability and vice versa. We estimated constancy stability as the fluctuation index (FI) of the time series (Dey and Joshi 2006) which is given as

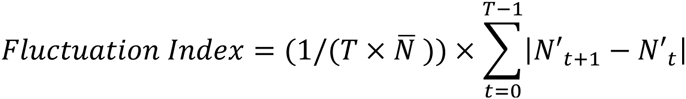

where *N*^′^_t_ denotes the breeding population size in *t*^th^ generation and 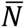 denotes average population size over *T* generations. Thus, higher values of FI suggest lower constancy and vice-versa.

Persistence stability of a population is a measure of its probability of extinction (Grimm and Wissel 1997) and was quantified as the proportion of generations in which a population went extinct, i.e.

*Extinction probability= Number of extinction events/Total number of generations* Clearly, higher extinction probability denotes lower persistence stability and vice versa. Since the control methods ensured that all extinctions were rescued, we scored the extinctions before the controls were implemented.

Effective population size (*N*_*e*_) is an indicator of how fast a population is expected to lose its genetic variation and thus, is a measure of its genetic stability (Hare et al. 2011). We estimated *N*_*e*_ as the harmonic mean of the time series (Allendorf and Luikart 2007), i.e.

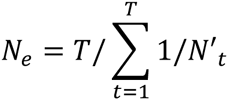

We also measured the average population size as the arithmetic of the population time series.

### 2.5 Effort magnitude and frequency

Following a previous study (Hilker and Westerhoff 2005) we estimated the effort magnitude, which is a proxy for the cost of implementation of the control methods. This quantity computes the number of individuals externally added or removed per generation and is given as:

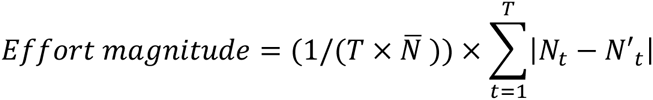

where *N*_*t*_ and *N*′_t_ are the population sizes before and after external perturbation in the *t*^*th*^ generation respectively. 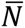 and *T* denote the average population size and length of the time series respectively (Sah et al. 2013).

Effort frequency was measured as the proportion of generations external perturbation was required *i.e.* the number of generations when perturbation was required divided by the total number of generations.

### 2.6 Statistical analyses

For statistical analyses, BLC and TOC were compared against the corresponding reference populations, C1 and C2, respectively. The data for the various indicators of stability (section 2.4) along with the effort magnitude and frequency (section 2.5) were subjected to separate one-way ANOVAs using STATISTICA^®^v5 (StatSoft. Inc., Tulsa, Oklahoma).We also computed the effect size of the difference between the means as Cohen’s *d* (Cohen 1988) using a freeware Effect Size Generator (Devilly 2004). Following standard recommendations (Cohen 1988), the value of effect size (*d*) was interpreted as large, medium and small when *d*>0.8, 0.8>*d*>0.5 and *d*<0.5 respectively.

### 2.7 Simulations

In order to test the generalizability of our empirical results, we performed biologically-realistic simulations using the Ricker model (Ricker 1954). Details of the simulations are mentioned in the supplementary section (Online Appendix A).

## 3. RESULTS

### 3.1 Both Limiter Control (BLC)

In the absence of control (*i.e.* for C1), population size distribution was observed to be positively skewed with a long right hand tail (figure 1A). When BLC was applied, the distribution became more symmetric with a higher mean and higher median population size (figure 1B). The absence of extreme values was reflected in terms of constancy and persistence. Compared to the C1 populations, the BLC populations had significantly reduced population fluctuation (figure 2A; F_1,8_=56.83, *p* = 0.00007, *d* = 4.77) as well as extinction probability (figure 2B; F_1,8_=26, *p*=0.0009, *d*= 3.23). Thus, both constancy and persistence stability were significantly enhanced and so were effective population size (figure 2C; F_1,8_=18.65, *p* = 0.003, *d* = 2.73) and average population size (figure2D; F_1,8_=23.21, *p*=0.001,*d =* 3.05). However, there was a steep cost to the attainment of this stability. BLC required significant effort magnitude (figure 2E, F_1,8_=574.56, *p* < 10^−8^,*d*= 15.16) and perturbations happened in almost every generation (figure 2F). Interestingly, although BLC involves both culling and restocking, here we found that culling events occurred more frequently than restocking ones (figure 3B) and are larger in magnitude (figure 3A). All the differences in means reported for BLC had large effect sizes (i.e. *d* > 0.8, see Table B1 for 95% CI around *d*).

### 3.2 Target Oriented Control (TOC)

The population size distribution of C2, like C1, had a long right hand tail (figure 4A). TOC prevented population sizes from taking extreme values, which made the distribution more symmetric with higher mean and median values (figure 4B). Thus, not surprisingly, TOC also increased both constancy and persistence stability by significantly decreasing fluctuation index (figure 5A; F_1,8_= 31.16, *p*= 0.001, *d*= 3.5) and extinction probability (figure 5B;F_1,8_= 32, *p*= 0.0005,*d*= 3.58) respectively. Interestingly, although TOC increased the effective population size of the controlled populations significantly (figure 5C; F_1,8_= 30.7, *p*= 0.001, *d*= 3.5), it had no effect on the average population size (figure 5D; F_1,8_=0.13, *p*=0.73, *d*= 0.23). Like BLC, the stability attained by TOC came at a high cost in terms of effort magnitude (figure 5E, F_1,8_= 921.16, *p* <10^−8^, *d*= 19.2) and frequency (figure 5F, F_1,8_= 591.4, *p* <10^−8^, *d*= 15.4). Unlike BLC though, in TOC, culling and restocking happened with almost similar frequency (figure 3D) and the effort magnitude for culling was only slightly greater than that for restocking (figure 3C). All the differences in means reported for TOC had large effect sizes (i.e. *d*> 0.8) except that for average population size, where it was found to be small (i.e. *d*< 0.5; see Table B2 for 95% CI around *d*).

### 3.3 Simulations

For both BLC and TOC, the outputs of the simulations using Ricker model under biologically realistic conditions followed the same trends as the experimental results (*cf* figure 2 and 5 with figure C1 and C2 respectively). The sole exception to this observation was the average population size for BLC. In experimental populations, BLC had significantly higher average population size than the unperturbed populations, whereas in simulations they have similar values (*cf* figure 2D and C1D).

## 4. DISCUSSION

Here we empirically studied the effects of two control methods in terms of inducing stability in unstable biological populations. However, it should be noted that this study does not concern itself with a direct comparison of the relative efficiency of these two methods. This is because it has already been shown numerically that most control methods are capable of inducing any desired level of stability with the correct choice of the value of the control parameter (Tung et al. 2014). Thus, comparisons of relative efficiency are possible only when both methods lead to the same level of stability, e.g. say a 50% enhancement of constancy as compared to the dynamics of the unperturbed populations (Tung et al. 2014). Clearly, in an experimental scenario, it is almost impossible to choose such parameter values *a priori* and therefore direct comparison of the efficiencies of the methods are improper. Hence, the goals of this study are to validate whether the methods can induce stability or not and to gain a biological understanding of the mechanisms by which these methods work. Thus, we prefer intuitive explanations based on how the control methods affect the population size distributions, instead of mathematically rigorous theorems and lemmas on how stability is attained.

**Figure 1.**
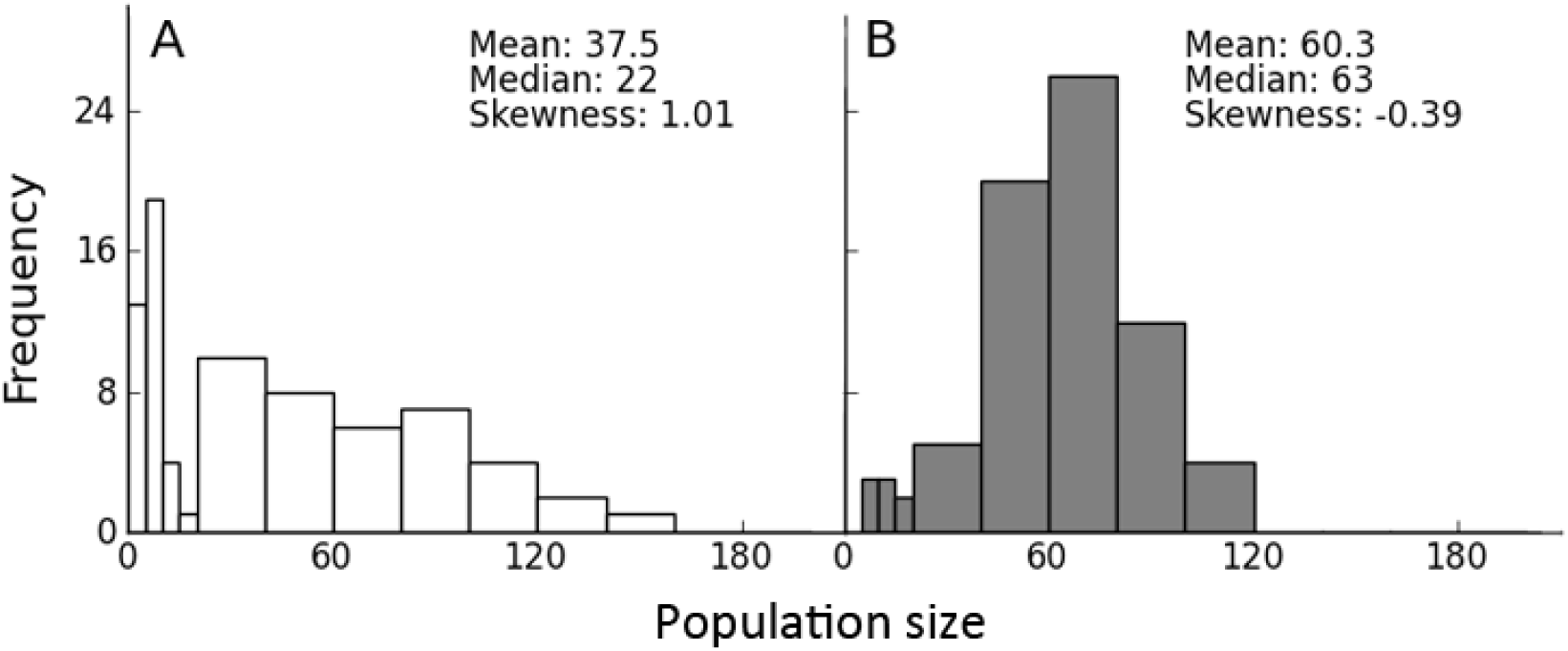
Population size distributions for unperturbed and BLC populations. **A**. Unperturbed (C1). **B.** BLC populations. Each distribution was plotted by pooling the data from all 5 populations over 14 generations. The bin size is 5 when the population size is in the range 0-20, and 20 otherwise. BLC made the distribution more symmetric and increased the measures of central tendencies (mean and median).

### 4.1 Both Limiter Control

*Population size distribution and constancy:* In the absence of any control, the distribution of population sizes had a long tail towards the right and about 49% of the values lay between 0-20 (figure 1A). As a result, the mean was much larger than the median which was also reflected as a high value for skewness (figure 1A). This kind of a distribution is a characteristic of populations undergoing high amplitude oscillations in sizes, which is known to be a feature of *Drosophila* populations subjected to a combination of low levels of larval nutrition and high levels of adult nutrition (Mueller 1988; Mueller and Huynh 1994). This is because high adult nutrition boosts the fecundity of the files which allows them to overcome the negative effects of density-dependence on fecundity at high population densities in a given generation (say *t*). As a result, a large number of eggs are laid, which leads to an overcrowding in the larval population of the next generation (*t+1*). Now, if the amount of larval nutrition available is less, the larval mortality is greatly increased, which in turn causes a crash in the adult numbers in generation *t+1*. However, the high fecundity of the flies ensures an immediate recovery from this trough in adult population size in the next generation (*t+2*), and thus the high-amplitude cycles continue (Mueller and Joshi 2000). Not surprisingly, the constancy stability of such dynamics is low (Mueller and Huynh 1994; Mueller and Joshi 2000; Sheeba and Joshi 1998).

When BLC was applied, the distribution became more symmetric as the right hand tail was curtailed and only 10.6% of the values lay in the range of 0-20 (figure 1B). Consequently, the mean came closer to the median and the value of the skewness was reduced (figure 1B). There were two reasons for this. Firstly, by definition, BLC ensured culling of individuals when the population size was above the upper threshold. Secondly, the presence of an upper threshold prevented over-crowding in the adult stage. This reduced the number of eggs laid in the next generation, ensuring lower egg-to-adult mortality and thus reducing the amplitude of population crashes. This is the biological equivalent of truncating the stock-recruitment curve (Hilker and Westerhoff 2005) and is known to reduce population size variability (Fryxell et al. 2005; Tung et al. 2015). Thus, not surprisingly, the fluctuation index of the BLC populations was found to be significantly lower than the corresponding unperturbed ones (figure 2A).However, this enhanced constancy came at a cost.

**Figure 2.**
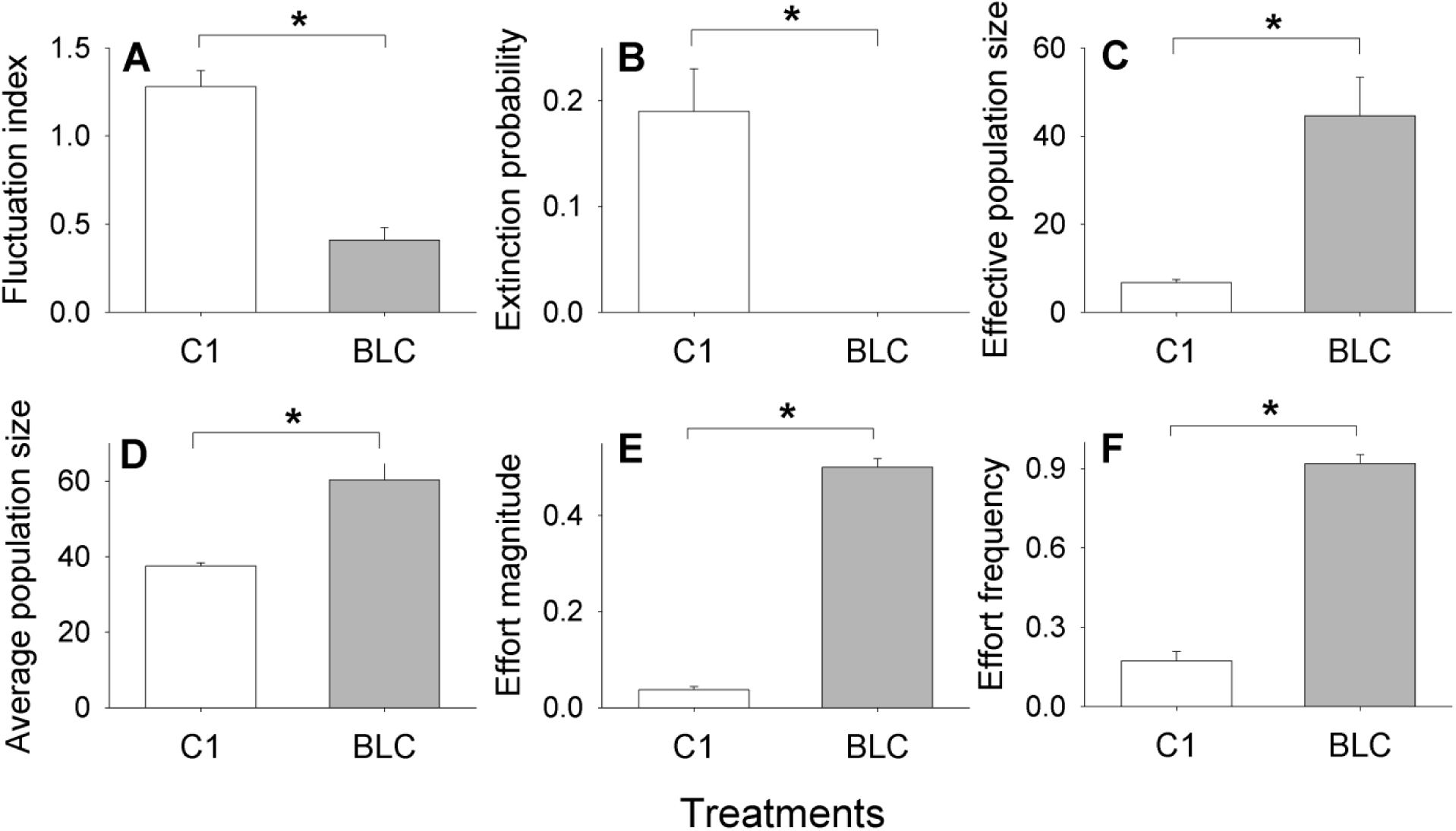
Empirical results for the effects of BLC. C1 represents corresponding unperturbed populations. Each bar represents a mean over 5 replicate populations. Error bars denote standard error around the mean and * denotes statistical significance (*p*< 0.05). BLC decreased **A. Fluctuation index** and **B. Extinction probability** significantly. It increased **C. Effective population size** and **D. Average population size** although at the cost of significantly high **E. Effort magnitude** and **F. Effort frequency**. See text for possible explanations.

*Effort magnitude and frequency:* Implementation of BLC required interventions in almost 90% of the generations (figure 2F) to the tune of ~30 individuals per generation (figure 2E). This implies that the effort required for BLC is likely to be prohibitively expensive for any real world application. Interestingly, although BLC provides for both culling and restocking, our data shows that ~99% of the total effort magnitude (figure 3A) and ~91% of the effort frequency (figure 3B) involved culling. This suggests that, in practise, it is the culling part of BLC which is likely to have a more significant impact on the dynamics. This is consistent with a recent empirical study (Tung et al. 2015) which shows that culling to a fixed upper threshold, *aka* Upper Limiter Control or ULC (Hilker and Westerhoff 2005), is capable of significant reduction in fluctuation in population sizes by itself. Thus, it is tempting to think of the restocking part of BLC as superfluous in terms of affecting the stability properties of populations. However, this simple line of reasoning was found to be erroneous, when we examined other kinds of stability.

**Figure 3.**
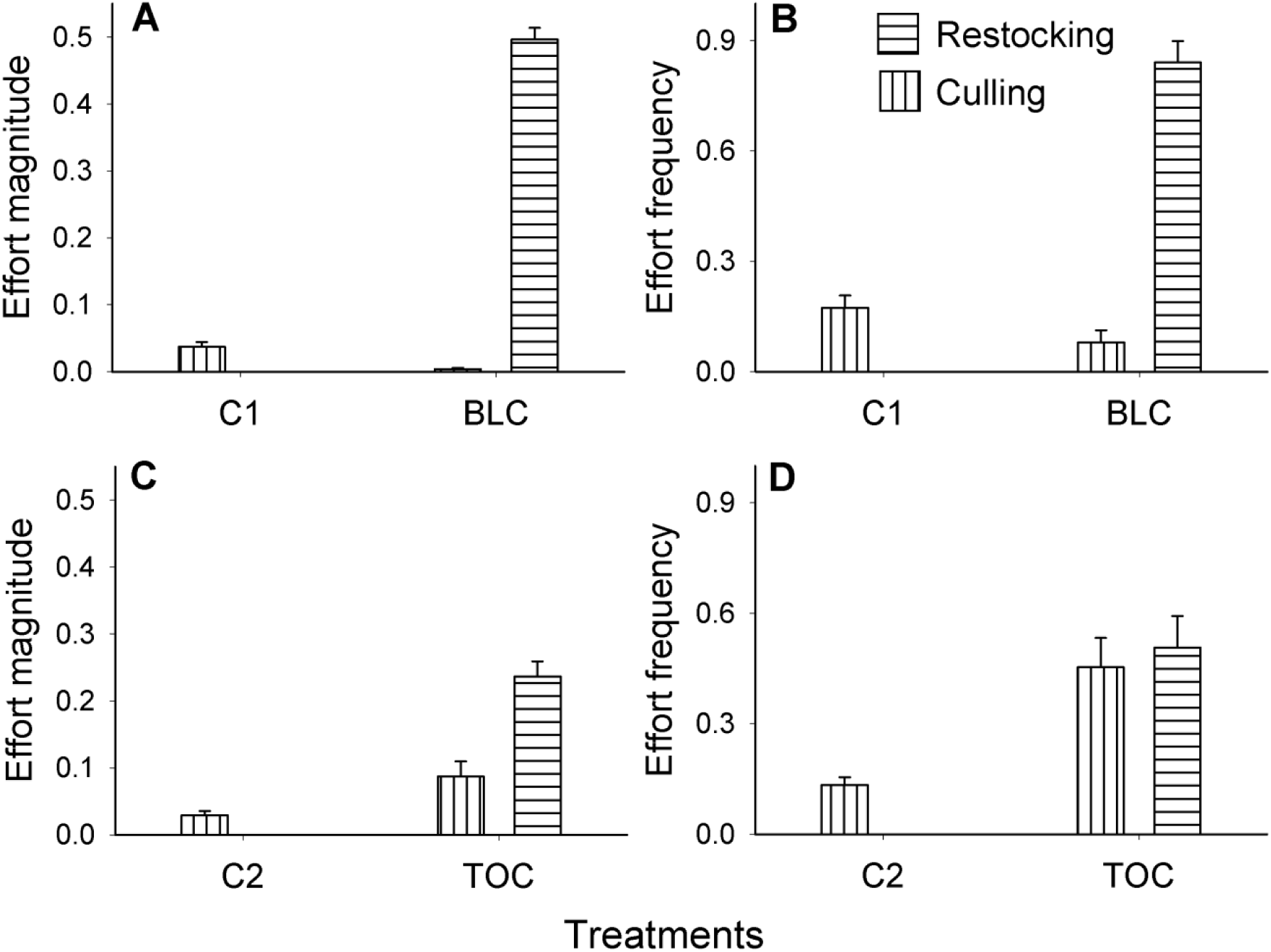
Magnitude and frequency of culling and restocking perturbation. Each bar represents a mean over 5 replicate populations and error bars denote standard error around the mean. C1 and C2 represent the unperturbed populations for BLC and TOC respectively. BLC mainly incurred culling perturbation w.r.t. both **A. Effort magnitude and B. Effort frequency**.TOC incurred more culling for. **C.Effort magnitude**, but similar amount of **D. Effort frequency** (see text for possible explanation and implications).

*Persistence stability:* An unperturbed *Drosophila* population can become extinct in two different ways. Firstly, as discussed above, large breeding population sizes in generation *t* can lead to extinction in the next generation (i.e. *t+1*), due to population crashes. Secondly, when the breeding population size is small, there is always a possibility of extinction in the next generation due to demographic stochasticity like all individuals being of the same sex or very low fecundity of the few available females etc (Lande 1988). In other words, both large and small breeding population sizes can lead to extinctions in the next generation. Although culling reduces the frequency of population crashes, it does not completely ameliorate it and therefore is unlikely to be a sufficient condition to increase persistence. This line of reasoning is supported by recent experimental results that showed that although ULC could cause some reduction in extinction probability, the effect was not statistically significant (Tung et al. 2015). BLC ameliorates this problem by putting a threshold to the lower values that the population can attain, and thus was more successful in promoting persistence (figure 2B). In fact, no extinctions happened in any of the 5 replicates that were controlled using BLC, which is also consistent with past theoretical studies (Tung et al. 2014).

*Effective and average population size:* A fluctuating population tends to lose genetic variability whenever it experiences population crashes (Allendorf and Luikart 2007). This can in turn lead to inbreeding-like effects, thus increasing the probability of extinction (Bijlsma et al. 2000). We quantified the effects of BLC on the genetic stability by estimating the effective population size (*N*_*e*_) as the harmonic mean of the population time series which is sensitive to low population numbers. BLC was found to significantly increase the *N*_*e*_ by about 3 times (figure 2C). Interestingly, a previous study that employed an upper threshold of 10 individuals (*i.e.* the same as the upper threshold of our BLC) failed to find a significant increase in *N*_*e*_ (Tung et al. 2015). This was primarily because there was a large variation in terms of the change in *N*_*e*_ which is in turn consistent with the observation that culling reduces the frequency of crashes in *Drosophila* populations, but does not ameliorate them totally. As BLC also ensures that the lower population size never goes below a pre-determined threshold, it completely rules out population crashes, thus causing a significant increase in *N*_*e*_.

Somewhat counter intuitively though, BLC also increased the average population size (figure 2D). Earlier theoretical studies have indicated that populations whose distributions have a long right tail (figure 1A) do not show an increase in average population size upon culling (Hilker and Westerhoff 2005). This prediction has also been verified by a recent study which employed the same level of culling as our current experiments (Tung et al. 2015). The only difference between the ULC treatment of the previous study (Tung et al. 2015) and our experiments is the restocking applied at very low magnitude (figure 3A) and frequency (figure 3B). However, this is insufficient to explain the observed increase in average population size as, unlike the harmonic mean, the arithmetic mean is not affected too much by low values in the population time series. Thus, it is not clear at this point as to why we observed an increase in the average population size in our experiments.

*Simulations:* Our simulations were able to capture all the trends of the empirical data (*cf* figure 2 with figure C1) except the one about average population size (figure C1D). Our simulation results predict no increase in average population size, which is consistent with earlier theoretical (Hilker and Westerhoff 2005) and empirical (Tung et al. 2015) studies on the effects of culling.

To summarize, at the level studied in this experiment, BLC enhanced all aspects of stability (constancy, persistence, effective population size) and average population size of unstable *Drosophila* population at the cost of very high effort.

### 4.2 Target Oriented Control (TOC)

*Population size distribution and constancy stability:* TOC is a two parameter control method in which a population is perturbed (i.e. culled or restocked) towards a pre-determined threshold (Dattani et al. 2011). For this experiment, the threshold was fixed at 30 (see methods for rationale) and each generation, 70% of the difference between the threshold and the present population size was either culled or restocked when the population size was greater or lesser than 30 respectively. Consequently, like BLC, TOC was also able to reduce the frequency of extreme values in the population size distribution (figure 4B). However, in the absence of fixed upper or lower thresholds, TOC was less effective in ameliorating the extreme values in the distribution (i.e. population booms and crashes) than BLC. This was reflected in the difference between the mean and the median and the higher value of skewness for TOC (*cf* figure 1B and figure 4B).The presence of greater number of extreme values might also be a result of the intentionally induced experimental noise, which was incorporated to put TOC to a more stringent test. Nevertheless, TOC was able to cause a significant reduction in fluctuation index (figure 5A).

**Figure 4.**
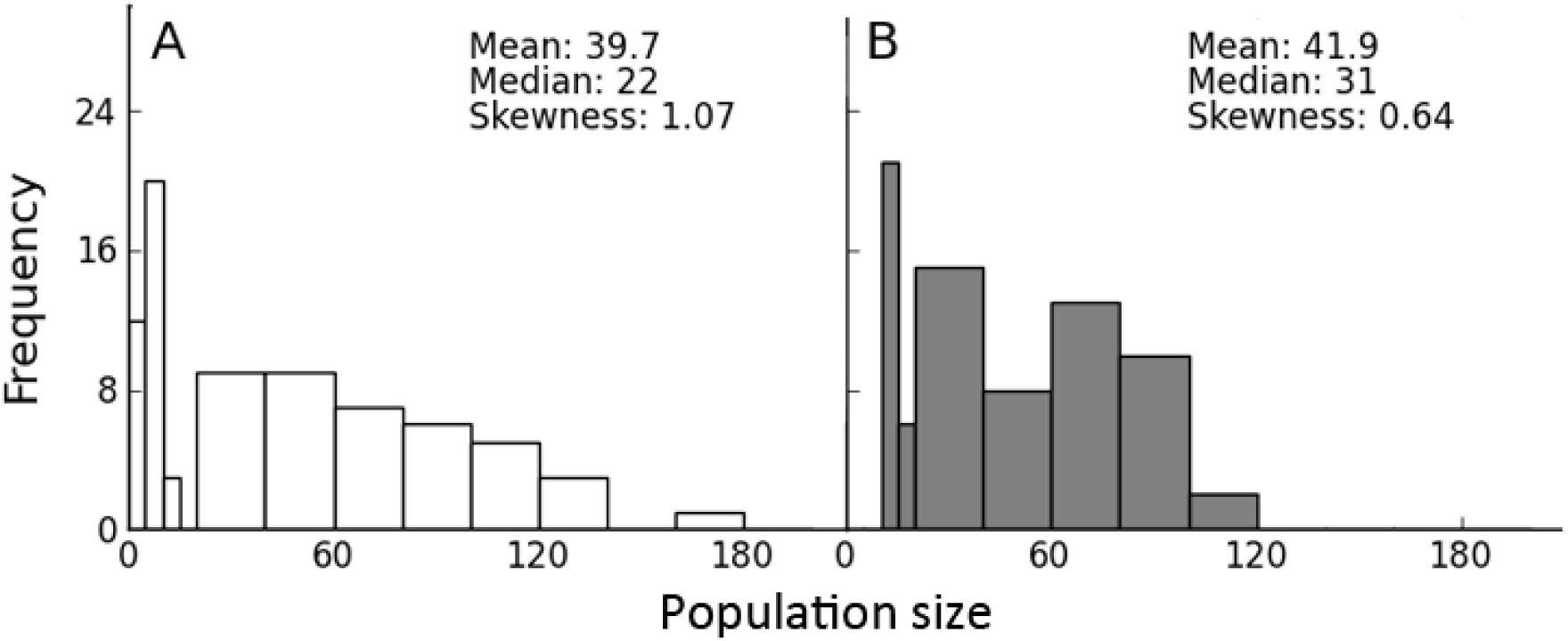
Population size distributions of the unperturbed and TOC populations. **A.** Unperturbed (C2). **B.** TOC populations. Each distribution was plotted by pooling the data from all 5 populations over 14 generations. The bin size is 5 when the population size is in the range 0-20, and 20 otherwise. TOC reduced the skewness and overall range of the population size distribution.

*Effort magnitude and frequency:* By its very design, application of TOC is expected to require culling or restocking except for the rare cases when population size would be exactly equal to the target value. Therefore, the effort frequency was found to be close to 1 (figure 5F). More crucially, there was a qualitative difference in terms of the pattern of the effort magnitude.

Unlike BLC, which consisted chiefly of culling (figure 3A, 3B), both culling and restocking were almost equally represented in the implementation of TOC in terms of magnitude (figure3C) as well as frequency (figure 3D). This difference in the relative frequency of culling and restocking between the two methods can again be explained by the population size distributions. In BLC, culling to a fixed upper threshold (10 females in our experiment) reduces the frequency of population crashes and therefore, leads to few opportunities for restocking. This is reflected by the fact that there are few values towards the left in figure 1B. Hence, almost the entire effort in terms of both magnitude (figure 3A) and frequency (figure 3B) is devoted to culling. On the other hand, by definition, TOC requires restocking whenever the number of individuals is less than 30. Clearly, for the level investigated in this study, TOC is less capable of checking increases in population sizes than BLC (*cf* figure 4B with figure 1B). This ensured that the restocking condition for TOC was met more times during the course of the experiment than the corresponding restocking condition for BLC (less than 4 females). Since TOC was relatively less effective in reducing population crashes, this line of reasoning leads to the prediction that TOC would be somewhat less effective than BLC in promoting persistence.

**Figure 5.**
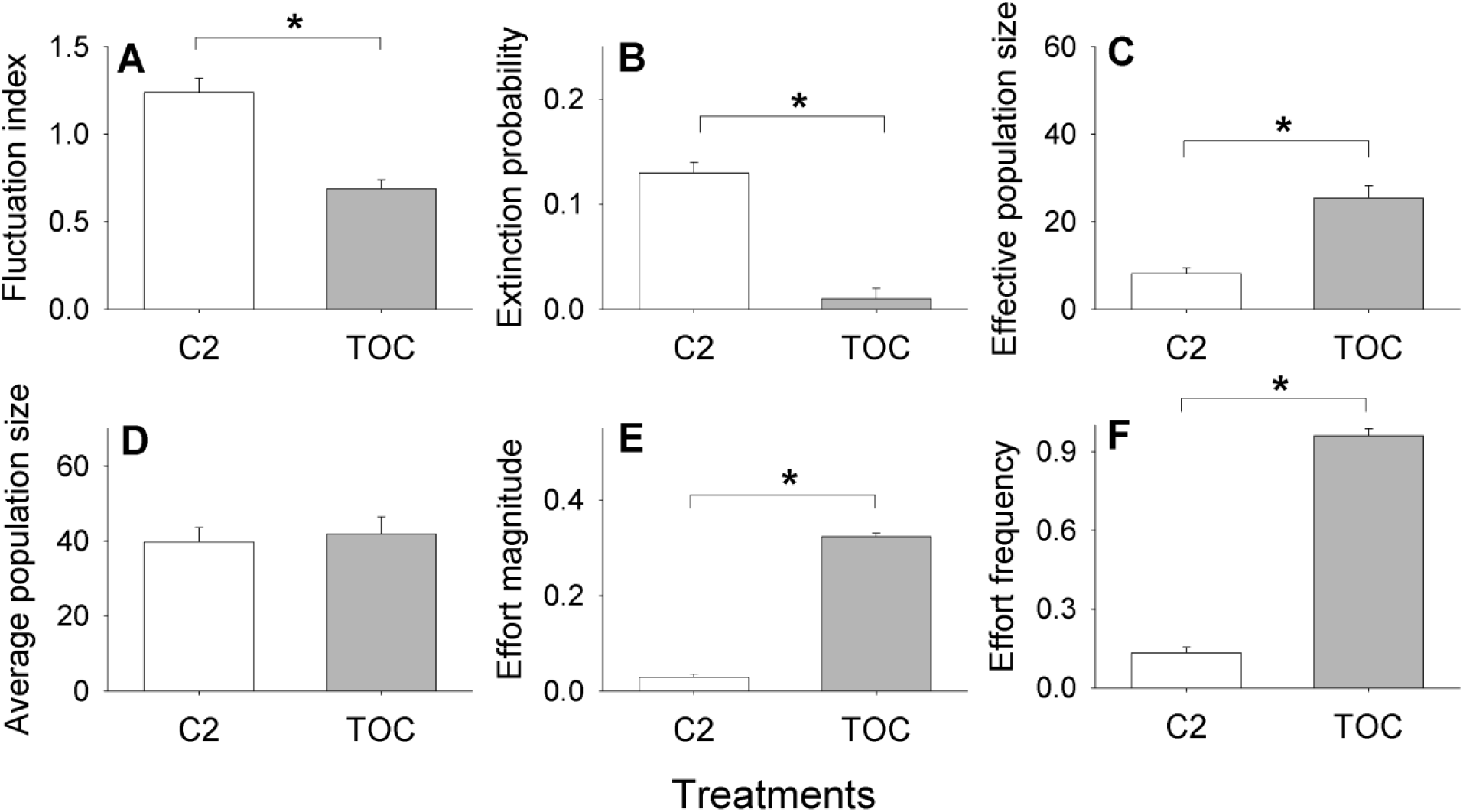
Empirical results for the effects of TOC. C2 represents the unperturbed populations. Each bar represents a mean over 5 replicate populations. Error bars denote standard error around the mean and * denotes *p*< 0.05. TOC decreased **A. Fluctuation index** and **B. Extinction probability** and increased **C. Effective population size** significantly, although it did not alter **D. Average population size.** TOC incurred significantly high **E. Effort magnitude** and **F. Effort frequency**. See text for possible explanations.

*Persistence stability, N_e_ and average population size:* TOC was capable of significantly reducing the extinction probability of the populations (figure 5B), although unlike BLC (figure 2B), it was not able to ameliorate extinctions altogether. However, as stated already, direct quantitative comparisons in terms of efficacy of the control methods are meaningless. More interesting is the observation that even with ~ 55% less effort than BLC (*cf* figure 2E and 5E), TOC was capable of a significant reduction in extinction probability (figure 5B). Similarly, the level of TOC investigated in this study was sufficient to induce a significant increase in effective population size (figure 5C) although the increase was not as much as that found in the BLC treatments. The reason for this can again be traced to the relatively large frequency of small sizes (thin bars in figure 4B) in the TOC populations as compared to the BLC populations (figure 1B) and the fact that *N*_*e*_ is more sensitive to small values. As expected from previous numerical studies (figure 5B of Dattani et al. 2011) and our own simulations (figure C2D), we found no difference in the average population size of the TOC populations (figure 5D).

Simulations: Our biologically realistic simulations were able to capture all the trends of the empirical data (*cf* figure 5 with figure C2), which ensures that these results are likely to be robust to noise and generalizable to other species.

In short, our study found that TOC increased constancy, persistence and *N*_*e*_ of unstable

*Drosophila* populations, but had no effect on average population size.

### 4.3 Caveats

Although we demonstrate that BLC and TOC can increase constancy, persistence and genetic stability of real biological populations, a number of caveats should be considered before extrapolating the results to field populations. Our experiments and simulations were performed on spatially unstructured, single species populations. However, most natural populations exist in multi-species communities and inhabit patchy habitats connected via migration. The dynamics of the population in such cases are expected to be more complex and it is not possible to infer how much of our results would be applicable to those scenarios. Moreover, while calculating the effort of implementation, we have ascribed similar weightage to both culling and restocking. But this may not be the situation in the wild, where depending on the context, culling might be easier than restocking or vice versa. Additionally, during restocking, individuals coming from outside may harbour new genetic variation and thereby alter the genetic makeup of the native population. Thus the effects of culling and restocking on the standing genetic variation may be fundamentally different, something that is ignored when *N*_*e*_ is used as a proxy for estimating how fast the population loses its variation. Furthermore, both these methods demand good estimates of the population sizes which might be an economically expensive affair to begin with. Finally, one must never lose sight of the biology of the controlled species as improper control can be disastrous for the ecosystems (Pyke 2008).

## Appendix A Description of simulations

We used the Ricker growth model (Ricker 1954) for simulating the dynamics of the unperturbed populations. This model is given as *N*_*t* + 1_ = *N*_*t*_\**exp*(*r*\*(1-*N*_*t*_/*K*)), where *r, K* and *N*_*t*_ denote intrinsic growth rate, carrying capacity and population size at time *t* respectively. The two control methods, Both Limiter Control (BLC) and Target Oriented Control (TOC) were imposed according to the following mathematical form (Tung et al. 2014):

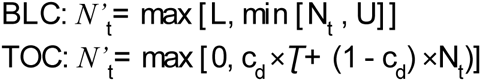

where *Nt* and *N*^′^_*t*_ are the population sizes before and after perturbation in the *t*^*th*^ generation. *N*_*t+1*_ = *f*(*N*^′^_*t*_), where *f* stands for Ricker model. *U* and *L* are the upper and lower thresholds of BLC while ***Ʈ*** and *c_d_* represent target and control intensity of TOC respectively. *min[]* and *max[]* stand for minimum and maximum operators.

*<u>Parameter values:</u>* We fit the Ricker model to the unperturbed experimental time-series (C1 and C2) and obtained a mean *r* and *K* of 2.6 and 34 respectively. These values of *r* and *K* were then used to simulate the dynamics of unperturbed and controlled populations. Matching the experimental values, upper (*U*) and lower (*L*) threshold for females in BLC were kept at 10 and 4 whereas the target (*Ʈ*) and control intensity (*c_d_*) for TOC were fixed at 30 and 0.7 respectively.

*<u>Stochasticity and lattice effect:</u>* Since noise can significantly influence the dynamics of perturbed populations (Dey and Joshi 2007), we incorporated noise to both the parameters, *r* and *K*, in each iteration. Following an earlier study (Tung et al. 2015) we picked the *r* and *K* values from uniform distributions of 2.6 ±0.5 and 34±15 respectively. Real organisms always come in integer numbers [lattice effect (Henson et al. 2001)], a constraint that can potentially affect the dynamics of the systems. We incorporated the lattice effect in our simulations by rounding off the number of organisms as well as the values of the perturbations in each generation to the nearest integer less than the actual value. The Ricker model, when initiated with a nonzero value, does not take zero-values and hence can never show extinctions. However, the same is not true for the integerized model. Therefore, we set reset rules similar to the experiment for our simulations: when the unperturbed populations went extinct, the population size was reset to 8. No resets were necessary for the simulations incorporating BLC and TOC as both control methods involved restocking steps.

The magnitudes of control to be applied were computed by assuming a 1:1 sex ratio. Thus, for BLC, the control method was implemented only when half the population size fell below 4 or exceeded 10. For TOC, the amount of perturbation was estimated by subtracting the population size before perturbation (*N*_*t*_) from the calculated post perturbation population size (*N*^′^_*t*_) and then only half of the perturbation value was actually realised in each generation.

To summarize, the sequence of steps in the simulation is given as:

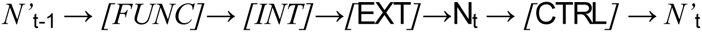

where *N*^′^_*t*_ and *N*_*t*_ are the population sizes after and before application of control in the *t*th generation, and *FUNC, INT, EXT* and *CTRL* represent the population recursion (here Ricker model), integerization, stochastic extinction and control (here BLC and TOC) functions respectively. The initial value for all simulations was set to 10, which was the same as the starting population size in the experiments. Each simulation was run for 100 iterations. From the resulting time series FI, extinction probability, average population size, effective population size, effort magnitude and effort frequency were computed over the resulting time series. All calculations, except extinction probability, were performed on the time series of the *N*^′^_*t*_ values. All figures (figure C1 and figure C2) represent means over 100 independent replicates for each treatment. This means that our simulations represent the dynamics of the populations over a longer time scale and larger number of replicates than what was performed in the experiments. This was important to study the long term behaviour of the populations and to ensure that our experimental results are not mere statistical artefacts. However, it is also important to ascertain whether we get back the same results if the simulations are repeated for the same number of replicates and number of generations as our experiments. Therefore, we also repeated all simulations for 14 generations and 5 replicates. Since there were no qualitative differences between the two cases, we chose to report the former set of simulations (i.e. 100 iterations and 100 replicates) here, as they represent the dynamics over a slightly longer timescale.

## Appendix B **Consolidated summary statistics**

**Table B1.** Summary statistics of the dynamics after applying BLC.

| **BLC** | Mean ±SE<br>for C1 | Mean ±SE<br>for BLC | ANOVA<br>F(1,8) | ANOVA<br>p value | Effect size<br>± 95% CI | Inference |
| --- | --- | --- | --- | --- | --- | --- |
| Fluctuation index | 1.28± 0.09 | 0.41± 0.07 | 56.83 | 0.00007 | 4.77± 2.43 | Large |
| Extinction probability | 0.19± 0.04 | 0 | 26 | 0.0009 | 3.23± 1.88 | Large |
| Effective population size | 6.8± 0.67 | 44.6± 8.7 | 18.65 | 0.003 | 2.73± 1.72 | Large |
| Average population size | 37.5± 0.84 | 60.3± 4.68 | 23.21 | 0.001 | 3.05± 1.82 | Large |
| Effort magnitude | 0.04± 0.01 | 0.5± 0.02 | 574.56 | $<10^{-8}$ | 15.16± 6.76 | Large |
| Effort frequency | 0.17± 0.03 | 0.92± 0.03 | 250.9 | $<10^{-6}$ | 10.02± 4.6 | Large |

\* C1 is unperturbed reference populations corresponding to BLC populations respectively. Cohen's $d$ statistic ( ± 95% confidence interval) was considered as the measure for effect size and interpreted as small, medium and large for $0.2 < d < 0.5$ , $0.5 < d < 0.8$ and $d > 0.8$ respectively.

**Table B2.** Summary statistics of the dynamics after applying TOC.

| **TOC** | Mean ±SE<br>for C2 | Mean ±SE<br>for TOC | ANOVA<br>F(1,8) | ANOVA<br>p value | Effect size<br>± 95% CI | Inference |
| --- | --- | --- | --- | --- | --- | --- |
| Fluctuation index | 1.24± 0.08 | 0.69± 0.05 | 31.16 | 0.001 | 3.5± 1.98 | Large |
| Extinction probability | 0.13± 0.01 | 0.01± 0.01 | 32 | 0.0005 | 3.58± 2 | Large |
| Effective population size | 8.08± 1.33 | 25.4± 2.83 | 30.7 | 0.001 | 3.5± 1.97 | Large |
| Average population size | 39.7± 3.94 | 41.9± 4.6 | 0.13 | 0.73 | 0.23± 1.24 | Small |
| Effort magnitude | 0.03± 0.006 | 0.32± 0.007 | 921.16 | $<10^{-8}$ | 19.2± 8.5 | Large |
| Effort frequency | 0.13± 0.02 | 0.96± 0.03 | 591.4 | $<10^{-8}$ | 15.4± 6.9 | Large |

\* C2 is unperturbed reference populations corresponding to TOC populations. Cohen's $d$ statistic ( ± 95% confidence interval) was considered as the measure for effect size and interpreted as small, medium and large for $0.2 < d < 0.5$ , $0.5 < d < 0.8$ and $d > 0.8$ respectively.

## Appendix C **Simulation Results**

**Figure C1.**
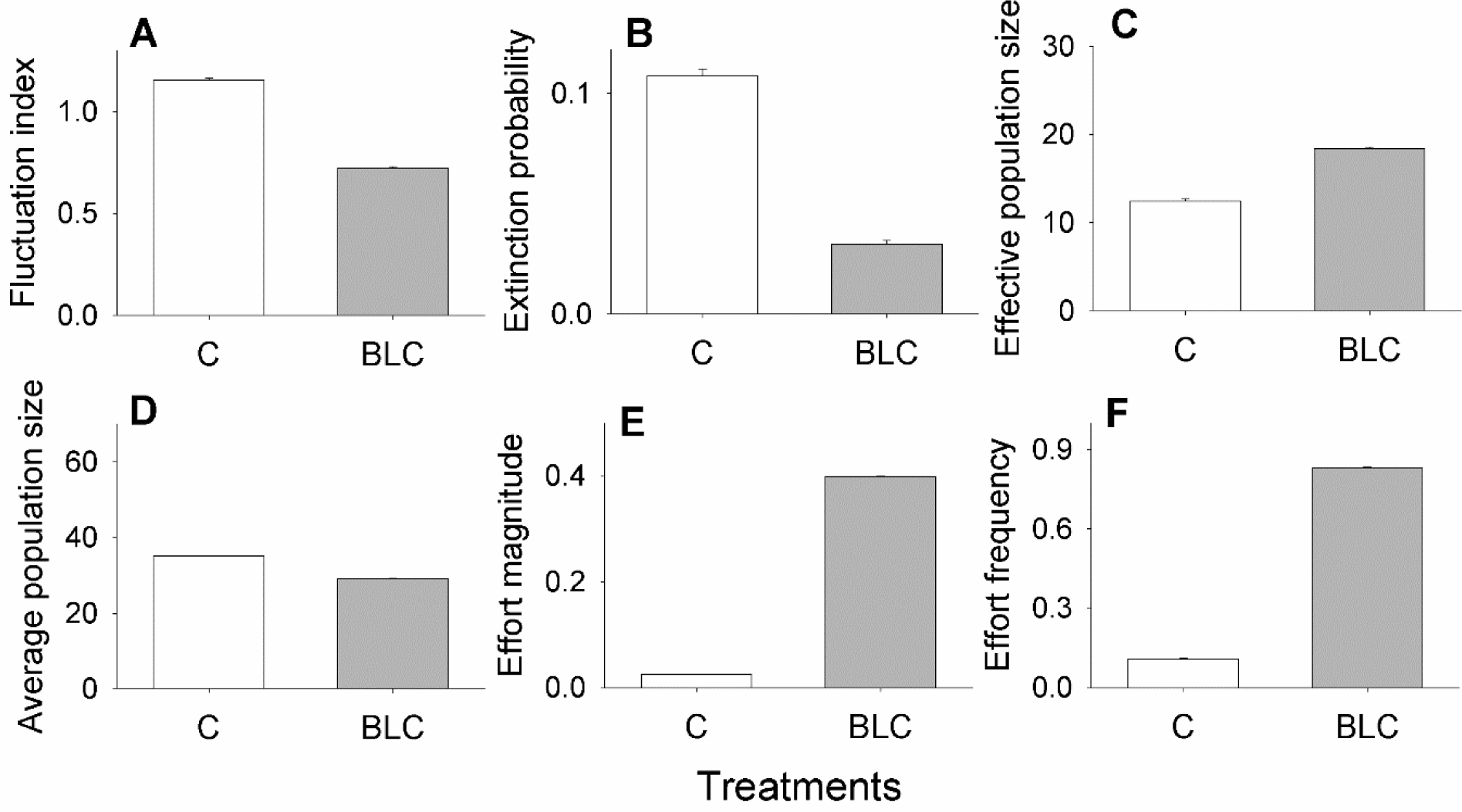
Simulation results for the effects of BLC. C represents unperturbed populations. Each bar represents a mean over 100 replicate populations and error bars denote standard error around the mean. **A. Fluctuation index**, **B. Extinction probability**, **C. Effective population size**, **D. Average population size**, **E. Effort magnitude** and **F. Effort frequency**.

**Figure C2.**
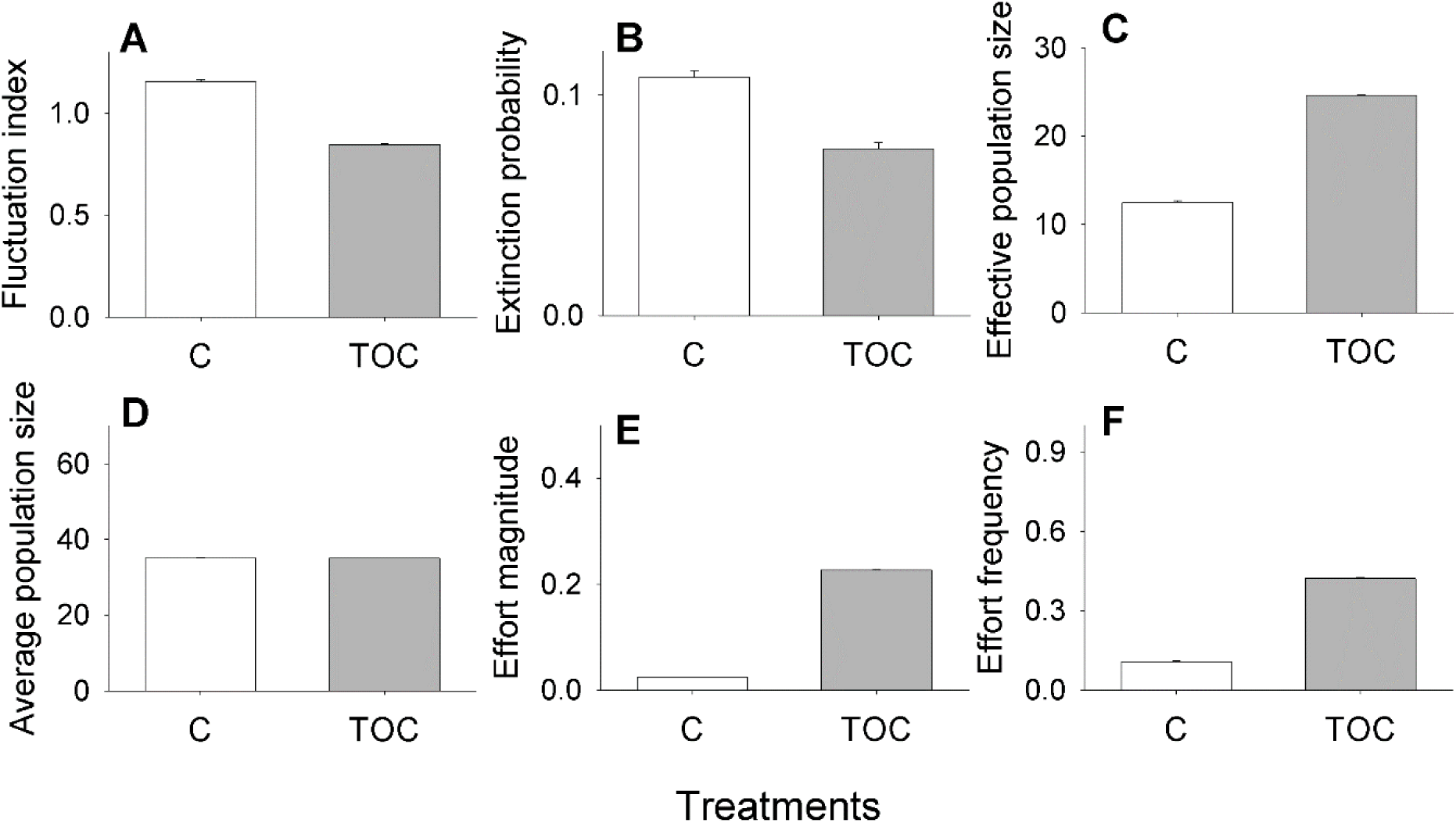
Simulation results for the effects of TOC. C represents unperturbed populations.Each bar represents a mean over 100 replicate populations and error bars denote standard error around the mean. **A. Fluctuation index**, **B. Extinction probability**, **C. Effective population size**, **D. Average population size**, **E. Effort magnitude** and **F. Effort frequency**.

